# Fast and furious: host aggression modulates behaviour of brood parasites

**DOI:** 10.1101/2020.04.27.063933

**Authors:** Václav Jelínek, Michal Šulc, Gabriela Štětková, Marcel Honza

## Abstract

Avian brood parasites pose a serious threat for hosts, substantially reducing their fitness which selects for the evolution of host defences. A classic example of a host frontline defence is mobbing which frequently includes contact attacking of brood parasites. Here, we investigated how the nest defence of a very aggressive great reed warbler (*Acrocephalus arundinaceus*) host influences the speed of egg-laying and egg-removing behaviour of its brood parasite – the common cuckoo (*Cuculus canorus*). We video-recorded 168 brood parasitic events at 102 active host nests and found that cuckoos avoided host mobbing in only 62% of cases. If hosts spotted the cuckoo at their nests, they almost always attacked it (in 91 of 104 cases), however, such attacks only rarely and temporarily prevented cuckoos from parasitizing (11 additional cases). When attacked, cuckoos parasitized host nests significantly faster and left them immediately. However, when not attacked, cuckoos frequently stayed at or near the nest suggesting that host aggression, rather than the risk of being spotted, influences the speed of brood parasitism in this species. Further, we found that cuckoos performed egg-removing behaviour in all parasitic events without regard to host aggression. As a result, cuckoos removed at least one egg during all brood parasitism events except those when an egg slipped from their beaks and fell back into the nest (in 9 of 73 cases). This indicates that egg-removing behaviour is not costly for the common cuckoo and is an essential part of its parasitism strategy, widening understanding of this currently unexplained behaviour.

## INTRODUCTION

Avian brood parasitism is an alternative reproductive strategy where parasites lay their eggs in the nests of hosts and transfer all parental care to them. The most extreme case is obligatory brood parasitism, occurring in about 1% of bird species, in which the parasites are unable to reproduce without their hosts (Yom-Tov 2001). During the evolutionary arms race between brood parasites and their hosts a lot of intriguing adaptations and counteradaptations have evolved. The most notable examples are egg mimicry of the common cuckoo (*Cuculus canorus*, Brooke and Davies 1988) or the cuckoo finch (*Anamalospiza imberbis*, Spottiswoode and Stevens 2011), egg rejection behaviour of their hosts (Wyllie 1981, Feeney et al. 2015), destroying of host eggs by cowbirds (*Molothrus* spp., Hoy and Ottow 1964) or killing of nestmates by nestlings of honeyguides (Indicatoridae, Spottiswoode and Koorevaar 2012).

One of the other remarkable adaptations of obligate brood parasites is their rapid egg-laying. While egg laying lasts from 20 to 100 min in non-parasitic species (Gill 2003, McMaster et al. 2004), brood parasites such as the common cuckoo (Moksnes et al. 2000, Andou et al. 2005), the brown-headed cowbird (*Molothrus ater*, McMaster et al. 2004) or the shiny cowbird (*Molothrus bonariensis*, Gloag et al. 2013) lay their eggs within seconds or even in a second like in a case of the great spotted cuckoo (*Clamator glandarius*, Soler et al. 2014). It has been suggested that rapid egg-laying has evolved to avoid being spotted by hosts that could use this information when defending against parasitism. Indeed, it was found that several host species react to the presence of stuffed or model brood parasite by increasing the rejection rate of model parasitic eggs (Davies and Brooke 1988, Moksnes et al. 1993, Bártol et al. 2012, Feeney et al. 2015, Samaš et al. 2016). Brood parasites could also minimize the time they spent at host nests to simply avoid or at least minimize the impact of host aggression as the less time they spend at the host nest the fewer attacks they suffer.

The role of host aggressivity has mostly been studied from the host perspective to show their ability to protect themselves against brood parasitism. However, it seems that host aggression against brood parasites is rather ineffective because even the most aggressive hosts suffer similar parasitism compared to less aggressive ones (Gill et al. 1997, Olendorf and Robinson 2000, Jelínek et al. 1994 but see Fiorini et al. 2009). Moreover, it has been shown that although hosts are sometimes capable of chasing the brood parasite away from their nests (Neudorf, and Sealy 1994, Webster 1994, Ellison and Sealy 2007, Gloag et al. 2013), they are not able to fend them off permanently and brood parasites finally succeed in almost all nests they choose (Neudorf, and Sealy 1994, Gloag et al. 2013, Soler et al. 2014).

From the parasite point of view, host aggressivity may be an important problem especially when parasites face relatively large and very aggressive hosts which are even able to kill them, such as great reed warblers parasitized by common cuckoos (Molnár 1944, Janisch 1948–51, Mérő and Žuljević 2014, Šulc et al. 2020) or chalk-browed mockingbirds (*Mimus saturninus*) parasitized by shiny cowbirds (*Molothrus bonariensis*, Gloag et al. 2013). Therefore, it should be beneficial for brood parasites to be as secretive as possible to avoid being noticed by hosts and when spotted to be as quick as possible to reduce potential injuries and the risk of death.

Here, we investigated how mobbing behaviour of one the most aggressive host species – the great reed warbler (hereafter GRW) - influences the behaviour of the common cuckoo (hereafter cuckoo) brood parasite. We predicted that cuckoos should shorten the time they spent at host nests when they are spotted or under host attack. We also expected that cuckoos should always leave the host nest immediately after laying to minimize the chance that hosts spot them because hosts may use this information as a cue for subsequent egg rejection (Davies and Brooke 1988, Moksnes et al. 1993, Bártol et al. 2012, Feeney et al. 2015, Samaš et al. 2016). Moreover, despite the direct benefits of fast brood parasitism, many parasite species hinder themselves with additional activities that apparently prolong the whole event; cowbirds peck and break host eggs to reduce the clutch size in host nest (Astié and Reboreda 2006) and cuckoos, including the common cuckoo, frequently remove one or more hosts eggs which they subsequently eat (Wyllie 1981). Therefore, we predicted that the time cuckoo females spend at empty host nests (where cuckoo cannot remove any egg) should be shorter than at nests containing eggs. Finally, we predicted that cuckoo females should cease egg removal when they are under host attack and instead quickly escape from aggressive hosts.

## METHODS

### Data collecting

The data were collected during the breeding seasons 2016, 2018 and 2019 in two adjacent fishpond areas between Hodonžn (48°51′N 17°07′E) and Mutěnice (48°54′N 17°02′E) in South Moravia, Czech Republic. The GRW population was individually marked and numbered 60, 45 and 65 breeding pairs in respective years and suffered a high rate of brood parasitism (73.7 % in 2016, 87.5 % in 2018 and 61.3 % in 2019). The majority of nests were found on the basis of regular mapping of male territories and checking for male mating status (Bensch 1996), the rest by systematically searching the littoral vegetation during nest building or early egg-laying stage and were checked typically daily until the clutch completion. Nests were checked less often (approximately every four days) during the incubation period except parasitized nests which were checked daily for five days following the clutch completion to determine the host response to parasitic eggs.

### Video-recording of nests

During all three breeding seasons we continually filmed the majority of great reed warbler nests at the study site during the egg-laying stage. The video-recording set-up comprised either a rear-view camera Carmedien STO-IR (Carmedien, Tschernitz, Germany) with IR illumination or custom HD cameras with inbuilt recorders without IR illumination and miniature digital videorecorder Mini DVR CH-HD0065 (Shenzhen Chu-Tech Co. Ltd, China) placed in a water-resistant box and powered by 12 V / 100 Ah gel batteries. The capacity of batteries was sufficient for about 7 days of continual recording, so in the majority of cases we did not need to change them during the whole recording period. All equipment was properly camouflaged. Batteries were placed on the bank far from nests. Water resistant boxes with recorders were hung on reeds and covered by masked cloths two to five metres from nests. Cameras were placed on thin aluminium poles camouflaged by cut reed stems and leaves fastened together by jute string. The distance between cameras and nest varied from 20 cm to two meters depending on the density of the reed (the denser the reed, the closer the camera was placed).

The set-up was installed mostly during the day preceding the day when the first GRW egg should be laid. The GRW nests are built exclusively by females and their construction lasts from three to four days ending when the nest cup is smooth and well-shaped. After that there is typically a one-day pause, during which females attend nests only occasionally to make some final adjustments of nest cups (Kluyver 1955, and our personal observations based on video recordings of nest building). Installing cameras at this time enhances the chance to film cuckoos attending nests and at the same time lowering the chance that warblers desert their nest because of our interference. This is of special importance as the installation of all equipment lasted from 30 minutes to one hour depending on the attributes of each nest site. As a result, we filmed egg-laying stage in 72 of 84 nests in 2016, 64 of 72 nests in 2018 and 58 of 75 nests in 2019. Of these 194 nests, 151 were filmed from the nest building stage, 33 from the day when one fresh egg was laid and 10 later during the egg-laying stage. Of 37 remaining nests which we did not film, 18 nests were found during incubation, three in the day when penultimate egg was laid, two with nestlings, three were found already deserted and empty (probably depredated) and eight during nest building but we did not film them due to logistical reasons (remote locations, temporary lack of equipment). Finally, we did not film three nests because we were aware that this female was extremely sensitive to disturbances in the vicinity of her nests. In total, we filmed 1225 nest/days. Filmed nests were visited mostly daily to check the nest content, adjust equipment and download the data (to change a memory card).

At the beginning of the 2016 breeding season we had problems with proper camouflage of our equipment and timing of its installation resulting in a higher frequency of nest desertion (13 cases including one caused by another reason). However, these problems were resolved later in the season and in future years, resulting in only one deserted nest due to insufficient camouflage in a second half of the 2016 breeding season and only seven in 2018 and 2019 together (three of them by the same sensitive female, see above). No other female deserted filmed nest twice and we filmed all their replacement nests which they start to build immediately after the desertion. In 2018 we performed a pilot experiment with additional nests similar to the experiments of Yang et al. (2017). As this experiment could substantially affect cuckoo female behaviour, we excluded all these nests from analyses (N = 19). We also excluded all deserted nests and events which took place after the nest desertion (as a response to brood parasitism). The final dataset thus comprised 168 brood parasitism events filmed at 102 nests.

### Analysis of video-recordings

To find all brood parasitism events (nest visits when cuckoo laid an egg) we watched all filmed material. For each event we recorded its length (defined as the time interval from the moment when the cuckoo sat on the nest rim to the moment of her departure), how many eggs were present in the nest (host’s and cuckoo’s together), how many eggs were taken by the cuckoo female, whether hosts were aware of the parasitizing cuckoo and whether they attacked it directly. Using the last two criteria we classified brood parasitism events into three categories: nests where the parents attacked cuckoo female (*aware and attacked*), nests where the parents were aware of parasitizing cuckoo but did not attacked it directly (*aware only*) and nests where the cuckoo female approached nest secretively and the parents did not know about her (*secretive approach*).

We were aware of the possibility that parents spotted approaching cuckoo but stayed out of the camera’s view and mobbed her without making contact. From this reason we performed additional analyses comparing the behaviour of hosts in events classified as *aware and attacked* and events classified as *secretive approach*. Firstly, we compared the behaviour of hosts during their first arrival after the brood parasitism event. GRWs which attacked parasitizing cuckoo usually behave very distinctly from the normal situation on their nest. They are noticeably excited, fluff feathers on the head, jump around the nest, look inside and sometimes peck egg(s). We encountered this behaviour in 85 of 91 nests where the parents attacked the cuckoo and in three of 62 nests originally classified as *secretive approach*. These three nests were, thus, reclassified as *aware only*. Secondly, we compared the time interval between the cuckoo’s departure and the first arrival of hosts on the nest between events classified as *aware and attacked, aware only* and *secretive approach* as parents should return faster after they mobbed the cuckoo. We found that hosts returned significantly sooner when they attacked the cuckoo (Wilcoxon test: W = 170, P < 0.001; *aware and attacked*: mean ± sd = 52 ± 85 s, range = 0 – 621 s, N = 91) or spotted the cuckoo but did not attack it (Wilcoxon test: W = 69, P < 0.001; *aware only*: mean ± sd = 129 ± 163 s, range = 8 – 623 s, N = 13) than when they were parasitized secretively (*secretive approach*: mean ± sd = 1100 ± 1268 s, range = 35 – 5539 s, N = 55, four cases when parents returned the next morning were excluded). Therefore, we believe that using above mentioned criteria enabled correct nest classification. Moreover, if we falsely classified some *aware only* nests as nests where the cuckoo approached the GRW nest secretively (*secretive approach*), it would bias our results in the opposite direction to our predictions. Hence, the classification used makes our conclusions more conservative.

### Statistical analysis

First, we investigated how the host aggression affects the speed of cuckoo brood parasitism. For this analysis we excluded five events where the cuckoos ate two eggs as these visits were extremely long (55, 59, 65, 93 and 114 seconds) making them clear outliers in our data (compare with data in Fig. 1). We also excluded 13 additional events where hosts were aware of the cuckoos but did not attack them to investigate solely the role of host aggression. We used generalized estimating equations (GEE; R package geepack, Yan 2002, Yan and Fine 2004, Hojsgaard et al. 2006) with a binomial error structure and exchangeable correlational structure as multiple brood parasitism was frequent in our data (43 of 93 nests in dataset were multiply parasitized) and the host response to the brood parasitism could depend on the host identity (their vigilance and nest guarding) and the attributes of the particular nest site (reed density, visibility for cuckoos etc., Jelínek et al. 2014). Since the response variable *brood parasitism length* had a right-skewed distribution with a steep increase and gradual decrease, we used GEE with gamma distribution. Variables *attack* (categorical with two levels: hosts did not know about the cuckoo and hosts attacked cuckoo directly) and *nest content* (categorical with two levels: empty nest and nest with egg) were included as predictors. We also included three potential confounding variables: *daytime, date* (both continuous) and a *year* (categorical). The model had this structure: `geeglm(brood_parasitism_length ∼ attack + nest_content + daytime + date + year, id` `= nestID, family = Gamma(link=log), corstr = “exchangeable”)`. We present the full model in the results.

**Figure 1:**
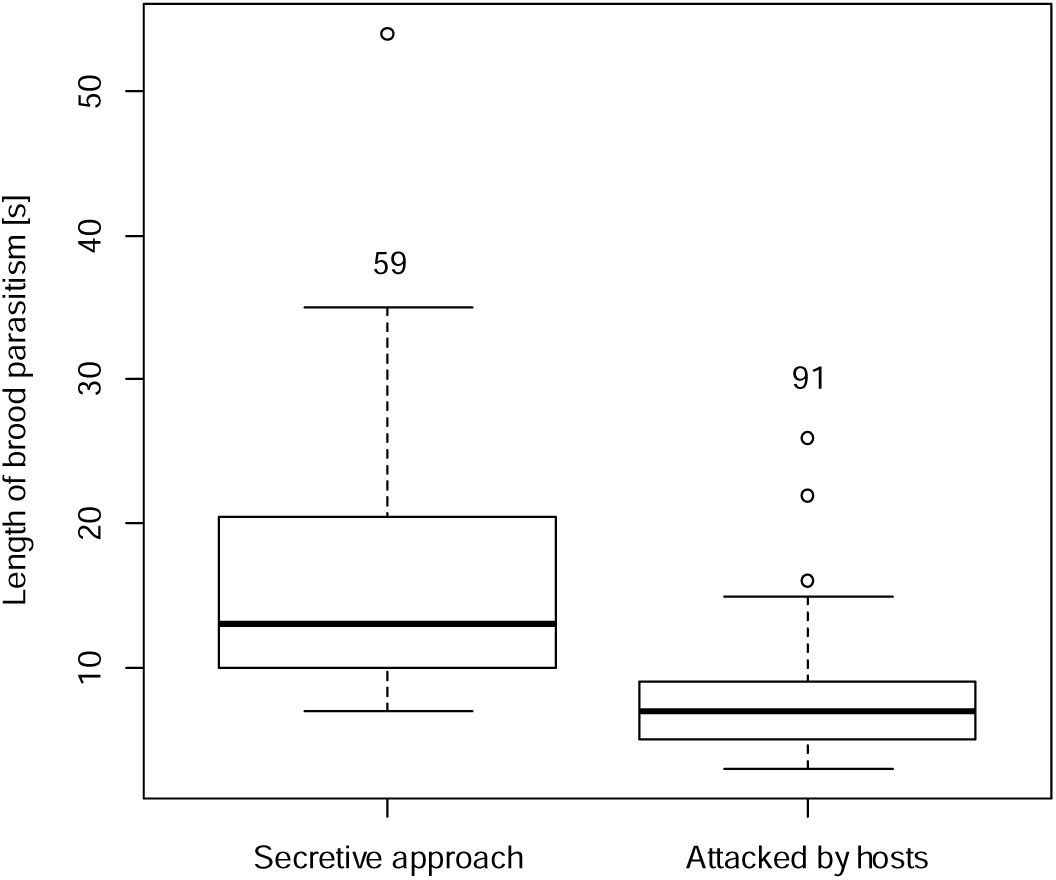
The length of the brood parasitism by the common cuckoo when it secretively approached the host nest or was attacked by great reed warbler hosts. Numbers above the boxplots represent sample sizes for each category. The bottom and top of the box represent the first and third quartiles and the bold band inside the box denotes the median. The whiskers indicate the lowest and highest data points that are within 1.5 the interquartile range, and data outside this range (circles) are regarded as potential outliers.

Second, the relationship between host aggression (*aware and attacked* vs. *secretive approach*) and the number of eggs which were taken by the cuckoo female during brood parasitism was investigated using a Chi-square test with continuity correction. From 168 filmed parasitism events, we omitted 13 events where hosts were aware of cuckoos but did not attack them and 28 events where cuckoos parasitized empty nests. The number of events thus lowered to 127 nests. As the number of eggs in the nest predetermines egg-taking options for cuckoos (cuckoo simply cannot remove more eggs than there are in the nest) and as cuckoos took between 0 and 2 eggs from our host nests, we made two comparisons. In the first comparison we used all 127 nests to test whether a cuckoo omits egg removal when she is attacked. In the second comparison, we used only events where host nests contained at least two eggs and cuckoos took at least one egg during brood parasitism (N = 77) to test whether cuckoos took more eggs when they are not attacked by hosts. All statistical analyses were performed in R 3.4.4 (R Core Team 2018).

## RESULTS

We video-recorded that hosts spotted the parasitizing cuckoo female in 104 of 168 cases and in these cases, they almost always attacked her (in 91 of 104 cases). Despite the fact that hosts attacked cuckoos fiercely and at such a high rate, we observed successful defence of the nest in only 11 cases.

Cuckoo females responded to host attacks by parasitizing about twice as fast (Table 1, Fig. 1); they spent on average 15.8 s at host nests (N = 59, SD = 8.3, range = 7 – 54 s; VIDEO 1) when they approached it secretively but 7.7 s (N = 91, SD = 3.5, range = 4 – 26 s; VIDEO 2) when they were under attack. Moreover, after parasitism cuckoo females immediately left the host nests without hesitation when attacked (in all 91 cases) while they sometimes stayed at host nests (in 16 of 59 cases, mean ± SD = 9.2 ± 4.2 s; VIDEO 3) or in their vicinity (in 3 of 59 cases; VIDEO 4) if hosts were not around. Contrary to our prediction, the length of parasitism events did not differ between empty nests and nests that contained eggs (Table 1).

**Table 1:** The effect of host vigilance and aggressiveness (*attack* – cuckoo secretively approached host nest vs. cuckoo directly attacked by hosts) and *nest content* (empty nests vs. nests with eggs) on the length of common cuckoo brood parasitism at the great reed warbler nests. *Attack* and *nest content* are categorical variables with two levels, *year* is a categorical variable with three levels (reference categories: ^a^cuckoo secretively approached host nest, ^b^empty nest, ^c^year 2016). Full model is presented.

| | df | Chi<sup>2</sup> | P | Estimate ± SE |
| --- | --- | --- | --- | --- |
| Intercept | | | | 2.977 ± 0.256 |
| Attack<sup>a</sup> | 1 | 75.0 | <0.001 | -0.601 ± 0.076 |
| Nest content<sup>b</sup> | 1 | 1.4 | 0.23 | -0.169 ± 0.096 |
| Daytime | 1 | 3.3 | 0.07 | -0.027 ± 0.015 |
| Date | 1 | 2.9 | 0.08 | $0.007 \pm 0.004$ |
| Year<sup>c</sup> | 2 | 10.8 | 0.004 | $0.217 \pm 0.086$ (year 2018: $P = 0.01$) |
| | | | | $0.227 \pm 0.086$ (year 2019: $P = 0.01$) |

Cuckoos also did not cease egg removal while attacked by hosts; at nests containing eggs, cuckoos removed at least one egg during all (N=127) but eight brood parasitism events. During these eight cases where cuckoos did not remove any egg, they were always attacked by hosts (Table 2a, χ^2^ = 4.69, P = 0,02). Only rarely (in 5 of 77 cases) did cuckoos remove two eggs from the host nest (Table 2b). During these five events, all cuckoos approached the host nests secretively, took one egg from it and ate it either there or in the close vicinity of the nest. Then they returned, removed the second egg, laid their own egg and finally left the host nest (VIDEO 5). Therefore, it seems, that egg removal of more than one egg is possible only when the nest is not guarded by hosts (χ^2^ = 11.8, P < 0,001).

**Table 2:**
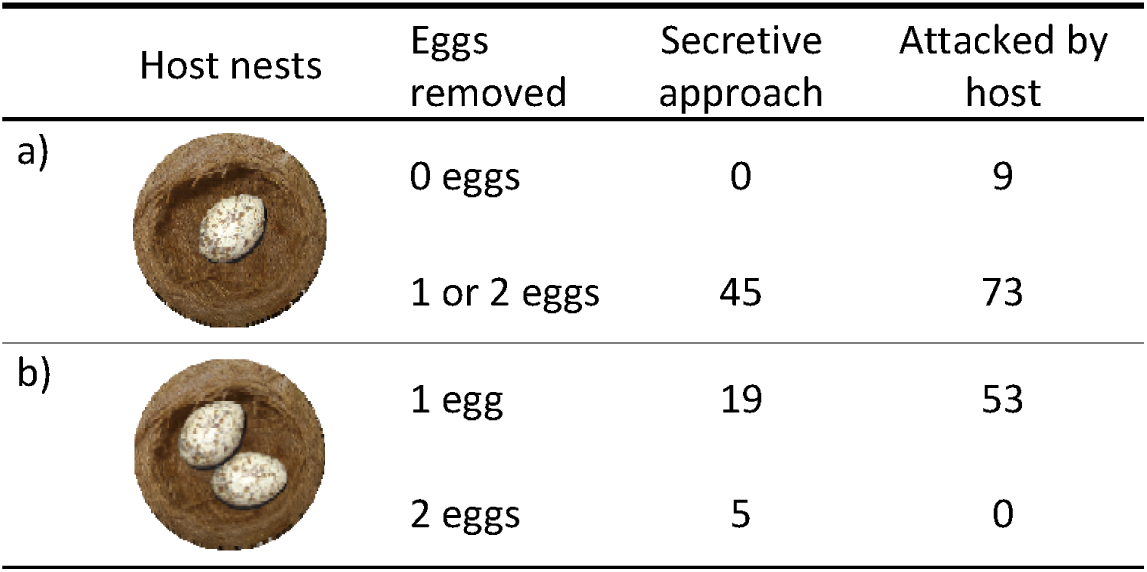
Number of eggs removed by cuckoo females during brood parasitism when they approached host nest secretively and when they were attacked by great reed warbler hosts – (a) at least one egg was present in host nest, (b) at least two eggs were present in host nest.

|  | Host nests | Eggs removed | Secretive approach | Attacked by host |
| --- | --- | --- | --- | --- |
| a) |  | 0 eggs | 0 | 9 |
|  |  | 1 or 2 eggs | 45 | 73 |
| b) |  | 1 egg | 19 | 53 |
|  |  | 2 eggs | 5 | 0 |

The same results were obtained for all analyses when the 13 *aware only* nests were added to dataset and treated as nests where the host attacked cuckoos (results not shown). The length of brood parasitism did not differ between the two groups of ‘aware’ nests (*aware and attacked* vs. *aware only*; Wilcoxon test: W = 632.5, P = 0.68).

## DISCUSSION

It has been proposed that common cuckoos should behave secretively during parasitism to lay their eggs without the host noticing (Davies and Brooke 1988, Moksnes and Røskaft 1989, Moksnes et al. 2000). Contrary to this general assumption, we found that the majority (62.0%) of GRW nests were parasitized when hosts were aware of the parasitizing cuckoo at their nests. Similar results were found for the reed warbler (*Acrocephalus scirpaceus*) by Moksnes et al. (2000) suggesting that at least these two hosts guard their nests heavily. Moreover, in most cases (87.5 %) where GRWs spotted the cuckoo female they mobbed her by direct attacks. These attacks were often very harsh; warblers were sitting on the cuckoos tearing their feathers out and pecking them in various places including the head (VIDEO 2).

Since GRWs attacked parasitic females vigorously, it is not surprising that we confirmed our first predictions; cuckoos sped up their parasitism when they were under host attack. We also found that cuckoos left host nests immediately after they laid but only when they were attacked. On the contrary, when hosts were not around, cuckoos laid an egg and then often stayed at nests or in their vicinity. This is surprising because cuckoos risked being spotted by hosts which has been shown to increase the chance of parasitic egg rejection (Davies and Brooke 1988, Moksnes and Røskaft 1989, Moksnes et al. 1993, Bártol et al. 2002). All these studies, however, were experimental, studying the reactions of hosts to stuffed or model cuckoos with an exposure time of at least five minutes (Davies and Brooke 1988, Moksnes and Røskaft 1989, Moksnes et al. 1993, Bártol et al. 2002). This is much longer than the duration of a real parasitism event which lasts on average 7.7 s in the GRW (this study) and 36.9 s in the reed warbler (Moksnes et al. 2000). Thus, it is possible that parasitism events that last just a few seconds seconds do not trigger a host rejection response at all, as hosts may consider cuckoos to be true nest predators that they quickly drove away and thus successfully saved their nest. The results of the only study (Moksnes et al. 2000) that tested the influence of the presence of real cuckoos at host nests on host rejection behaviour support the previously mentioned experimental studies. However, its results should be taken with caution due to the small sample size (N = 13 parasitized nests). Thus, further studies investigating the host reaction to cuckoos during real parasitism events are required to better understand the rejection behaviour of hosts.

Other brood parasites such as the great spotted cuckoo or the shiny cowbird also frequently expose themselves to attacks of larger and extremely aggressive hosts during brood parasitism (71% and 83% respectively) and these strikes could be even much more brutal that those executed by GRWs (Gloag et al. 2013, Soler et al. 2014). Similar to our study, these two parasites also spend less time on host nests when attacked by hosts showing that brood parasites at least try to lower the impact of host mobbing (Gloag et al. 2013, Soler et al. 2014). Cuckoos parasitizing reed warblers spend less time on host nests also in the study of Moksnes et al. (2000) but this relationship was not statistically significant probably because of the small sample size (N = 14) or perhaps because reed warblers do not usually attack the cuckoo directly (Čapek et al. 2010).

Interestingly, our observations do not support the hypothesis that egg-removing behaviour by parasitizing cuckoos prolongs the time they spend on host nests. We found that cuckoos spent a similar amount of time time when parasitizing empty nests (where they could not remove any egg) compared to parasitizing nests containing eggs. Moreover, we observed cuckoos removing eggs during all parasitic events and they did not omit this behaviour even when attacked by hosts. We recorded only 9 of 127 brood parasitism events where cuckoos did not remove an egg. However, after thorough inspection of video-recordings, in all these events, the cuckoos tried to remove the egg but failed to do it. The cuckoos tried to grasp an egg but were attacked by hosts so brutally that they dropped it back after they laid their own egg or did not manage to grasp it properly (VIDEO 6). After that, the cuckoos did not try to remove an egg again, probably because they could mistakenly take their own egg instead. Cuckoos also removed at least one host egg during all 14 brood parasitism events filmed by Moksnes et al. (2000) in reed warbler nests. Therefore, it seems that in the cuckoo the egg-removing behaviour must always precede the egg-laying. Even in empty nests, cuckoos searched for an egg, double checking whether there was truly nothing to take and laid its own egg only when they were sure that the nest was truly empty. On the contrary, in the nests that contained egg(s) cuckoos simply grabbed one of them and immediately laid their own egg (see VIDEO 7 for comparison). These two main components of cuckoo brood parasitism (egg removing and egg laying) follow each other so quickly and are usually so well executed that in an ideal case the whole brood parasitism event can take only four seconds. Interestingly, the egg laying frequently happens when the cuckoo has its head (with egg in beak) still inside the nest cup (VIDEO 8), reducing the delay caused by egg-removing behaviour to the minimum of several seconds.

Contrary to egg-removal, egg puncturing behaviour exhibited by the shiny cowbird really prolongs the time parasitic females spend on host nests. It has been shown that they need only 6.3 s for successful egg-laying, but on average 26 s when they peck host eggs. As a result, they get 17 blows on average to their head and body with beaks of their mockingbird hosts (Gloag et al. 2013) which suggests that egg-puncturing has similar importance for shiny cowbirds as egg-removing behaviour for cuckoos.

Besides 168 brood parasitism events we also filmed seven events in four nests where the hosts chased cuckoos away (VIDEO 9). In these cases, cuckoos withdrew immediately after hosts attacked them and did not attempt to approach the nest. Four other video-recordings from four nests showed cuckoos approaching host nest successfully but harsh attacks of GRWs prevented them from egg laying. Unfortunately for GRWs, these cases are probably very rare as six of these seven nests where successful defence occurred were subsequently parasitized. The last nest was parasitized three times before the successful defence was recorded. Then, the hosts subsequently deserted the nest. Thus, its is not possible to determine the reason why the fourth cuckoo female did not parasitize it. Similarly, poor success rate of host nest defence has been found by Gloag et al. (2013) where only in 17 of the 213 visits did the chalk-browed mockingbirds prevent shiny cowbirds from egg-laying.

To conclude, we show that even very harsh mobbing of GRWs does not save their nests from brood parasitism or even egg loss caused by egg-removing behaviour of cuckoo females. On the other hand, cuckoos react to host aggression and sped up the brood parasitism when attacked. This quickening mainly affects the egg-searching and egg-removing behaviour as the egg-laying alone lasts in all cases approximately two seconds. On the other hand, cuckoo females never avoid removing an egg and try to find an egg even in empty host nests. This suggests that the egg-removing behaviour is a more important part of the cuckoo’s brood parasitic strategy than was previously thought, although despite several proposed hypotheses and dozens of studies we can still only speculate about its adaptive reasons (reviewed in Šulc et al. 2016). Future studies focusing in detail on the process of brood parasitism in different species of evicting cuckoos can help to understand the adaptive significance of this peculiar behaviour.

## ACKNOWLEDGEMENTS

This work was supported by a project from the Czech Science Foundation (grant number 17-12262S). MŠ was funded by a Subsidy for research and mobility support of starting researchers of the Czech Academy of Sciences (MSM200931801). We thank V. Brlík, J. Koleček, K. Sosnovcová, R. Valterová-Poláková, S. McClelland, M. Cordall, B. Prudík, A. Aspinwall, F. Svoboda, M. Čapek, P. Procházka and M. Požgayová for their assistance in the field. We thank also to Anna E. Hughes for her English proof-read. Finally, we are obliged to the management of the Fish Farm Hodonín and local conservation authorities for permission to conduct the fieldwork. The study complied with the current laws and ethical guidelines of the Czech Republic.

## DATA AVAILABILITY STATEMENT

Data available in article supplementary material

## REFERENCES

Astié, A.A. & Reboreda, J.C. 2006. Costs of egg punctures and parasitism by shiny cowbirds (*Molothrus bonariensis*) at creamy-bellied thrush (*Turdus amaurochalinus*) nests. Auk 123: 23–32.

Andou, D., Nakamura, H., Oomori, S. & Higuchi, H. 2005. Characteristics of brood parasitism by common cuckoos on azure-winged magpies, as illustrated by video recordings. Ornithol Sci 4: 43–48.

Baltz, M.E. & Thompson, C.F. 1988. Successful incubation of experimentally enlarged clutches by house wrens. Wilson Bull 100: 70–79.

Bártol, I., Karcza, Z., Moskát, C., Røskaft, E. & Kisbenedek, T. 2002. Responses of great reed warblers *Acrocephalus arundinaceus* to experimental brood parasitism: the effects of a cuckoo *Cuculus canorus* dummy and egg mimicry. J Avian Biol 33: 420–425.

Bensch, S. 1996. Female mating status and reproductive success in the great reed warbler: is there a potential cost of polygyny that requires compensation? J Anim Ecol 65: 283–296.

Brooke, M.D. & Davies, N.B. 1988. Egg mimicry by cuckoos *Cuculus canorus* in relation to discrimination by hosts. Nature 335: 630–632.

Brooker, M.G. & Brooker, L.C. 1989. The comparative breeding behaviour of two sympatric cuckoos, Horsfield’s bronze-cuckoo *Chrysococcyx basalis* and the shining bronze-cuckoo *C. lucidus*, in Western Australia: a new model for the evolution of egg morphology and host specificity in avian brood parasites. Ibis 131: 528–547.

Brooker, M.G. & Brooker, L.C. 1991. Eggshell strength in cuckoos and cowbirds. Ibis 133: 406–413.

Čapek, M, Požgayová, M., Procházka, P. & Honza, M. 2010. Repeated presentations of the common cuckoo increase nest defense by the Eurasian reed warbler but do not induce it to make recognition errors. Condor 112: 763–769.

Chance, E.P. 1922. The Cuckoo’s Secret. London: Sidgwick and Jackson.

Davies, N.B. & Brooke, M. de L. 1988. Cuckoos versus reed warblers: adaptations and counteradaptations. Anim Behav 36: 262–284.

Davies, N.B. 2000. Cuckoos, cowbirds and other cheats. London: T & AD Poyser.

Ellison, K. & Sealy, S.G. 2007. Small hosts infrequently disrupt laying by brown-headed cowbirds and bronzed cowbirds. J Field Ornithol 78: 379–389.

Engstrand, S.M. & Bryant, D.M. 2002. A trade-off between clutch size and incubation efficiency in the Barn Swallow *Hirundo rustica*. Funct Ecol 16: 782–791.

Feeney, W.E., Troscianko, J., Langmore, N.E. & Spottiswoode, C.N. 2015. Evidence for aggressive mimicry in an adult brood parasitic bird, and generalized defences in its host. Proc R Soc B 282: 20150795.

Fiorini, V.D., Tuero, D.T. & Reboreda, J.C. (2009). Host behaviour and nest-site characteristics affect the likelihood of brood parasitism by shiny cowbirds on chalk-browed mockingbirds. Behaviour, 1387–1403.

Gärtner, K. 1981. Das Wegnehmen von Wirtsvogeleiern durch den Kuckuck (*Cuculus canorus*). Ornithol Mitt 33: 115–131.

Gill, S.A., Grieef, P.M., Staib, L.M. & Sealy, S.G. 1997. Does nest defence deter or facilitate cowbird parasitism? A test of the nesting cue hypothesis. Ethology 103: 56–71.

Gill Sharon A. 2003. Timing and duration of egg laying in duetting buff-breasted wrens. J Field Ornithol 74: 31–36.

Gloag, R., Fiorini, V.D., Reboreda, J.C. & Kacelnik, A. 2013. The wages of violence: mobbing by mockingbirds as a frontline defence against brood-parasitic cowbirds. Anim Behav 86: 1023–1029.

Grim, T., Rutila J., Cassey, P. & Hauber, M.E. 2009. The cost of virulence: an experimental study of egg eviction by brood parasitic chicks. Behav Ecol 20: 1138–1146.

Hamilton, W.J.I. & Orians, G.H. 1965. Evolution of brood parasitism in altricial birds. Condor 67: 361–382.

Hojsgaard, S., Halekoh, U. & Yan, J. 2006. The R package geepack for generalized estimating equations. J Stat Softw 15: 1–11.

Hoy, G. & Ottow, J. 1964. Biological and oological studies of the molothrine cowbirds (Icteridae) of Argentina. Auk 81: 186–203.

Jaatinen, K., Öst, M., Waldeck, P. & Andersson M. 2009. Clutch desertion in Barrow’s goldeneyes (*Bucephala islandica*) – effects of non-natal eggs, the environment and host female characteristics. Ann Zool Fennici 46: 350–360.

Janisch, M. 1948–51. Fight between *Cuculus c. canorus* L. – cuckoo – and *Acrocephalus a. arundinaceus* L. – great reed warbler. Aquila 55–58: 291.

Jelínek, V., Procházka, P., Požgayová, M. & Honza, M. 2014. Common Cuckoos *Cuculus canorus* change their nest-searching strategy according to the number of available host nests. Ibis 156: 189–197

Kluyver, H. N. (1955) Das Verhalten des Drosselrohrsängers, *Acrocephalus arundinaceus* (L.), am Brutplatz mit besonderer Berücksichtigung der Nestbautechnik und der Revierbehauptung. Ardea 43: 1–50.

Lerkelund, H.E., Moksnes, A., Røskaft, E. & Ringsby, T.H. 1993. An experimental test of optimal clutch size of the fieldfare; with a discussion on why brood parasites remove eggs when they parasitize a host species. Ornis Scand 24: 95–102.

McMaster, D.G. & Sealy, S.G. 1997. Host-egg removal by brown-headed cowbirds: A test of the host incubation limit hypothesis. Auk 114: 212–220.

McMaster, D.G., Neudorf, D.L.H., Sealy, S.G. & Pitcher, T.E. 2004. A comparative analysis of laying times in passerine birds. J Field Ornithol 75: 113–122.

Mérő, T.O. & Žuljević, A. 2014. Great reed warbler *Acrocephalus arundinaceus*. Acrocephalus 34: 130.

Moksnes, A. & Roskaft, E. 1989. Adaptations of meadow pipits to parasitism by the common cuckoo. Behav Ecol Sociobiol 24: 25–30.

Moksnes, A., Røskaft, E. & Korsnes, L. 1993. Rejection of cuckoo (*Cuculus canorus*) eggs by meadow pipits (*Anthus pratensis*). Behav Ecol 4: 120–127.

Moksnes, A., Røskaft, E., Hagen, L.G., Honza, M., Mørk, C. & Olsen, P.H. 2000. Common cuckoo *Cuculus canorus* and host behaviour at reed warbler *Acrocephalus scirpaceus* nests. Ibis 142: 247–258.

Molnár, B. 1944. The Cuckoo in the Hungarian plain. Aquila 51: 100–112.

Moskat, C. & Honza M. 2002. European cuckoo *Cuculus canorus* parasitism and host’s rejection behaviour in a heavily parasitized great reed warbler *Acrocephalus arundinaceus* population. Ibis 144: 614–622.

Neudorf, D.L. & Sealy, S.G. 1994. Sunrise nest attentiveness in cowbird hosts. Condor 96: 162–169.

Nord A, & Nilsson, J-Å. 2012. Context-dependent costs of incubation in the pied flycatcher. Anim Behav 84: 427–436.

Olendorf, R. & Robinson, S.K. 2000. Effectiveness of nest defence in the Acadian flycatcher *Empidonax virescens*. Ibis 142: 365–371.

Picman, J. & Pribil, S. 1997. Is greater eggshell density an alternative mechanism by which parasitic cuckoos increase the strength of their eggs? J Ornithol 138: 531–541.

R Core Team. 2018. R: A language and environment for statistical computing. R Foundation for Statistical Computing, Vienna, Austria.

Samaš, P., Rutila, J. & Grim, T. 2016. The common redstart as a suitable model to study cuckoo-host coevolution in a unique ecological context. BMC Evol Biol 16: 255

Scott, D., Weatherhead, P.J. & Ankney, C.D. 1992. Egg-eating by female brown-headed cowbirds. Condor 94: 579–584.

Sealy, S.G. 1992. Removal of yellow warbler eggs in association with cowbird parasitism. Condor 94: 40–54.

Soler, M., Pérez-Contreras, T. & de Neve, L. 2014. Great spotted cuckoos frequently lay their eggs while their magpie host is incubating. Ethology 120: 965–972.

Spottiswoode, C.N. & Stevens, M. 2011. How to evade a coevolving brood parasite: Egg discrimination versus egg variability as host defences. Proc R Soc B 278: 3566–3573.

Spottiswoode, C.N., & Koorevaar, J. 2012. A stab in the dark: chick killing by brood parasitic honeyguides. Biol Lett 8: 241–244.

Šulc, M., Procházka, P., Čapek, & Honza, M. 2016. Common cuckoo females are not choosy when removing an egg during parasitism. Behav Ecol 27: 1642–1649.

Šulc M. Štětková G. Procházka P. Požgayová M. Studecký J. & Honza M. 2020. Caught on camera: circumstantial evidence for fatal mobbing of an avian brood parasite by a host. J. Vertebr. Biol. 69: 20027.

Yan, J. 2002. Geepack: yet another package for generalized estimating equations. R-News 2/3: 12–14.

Yan, J. & Fine, J.P. 2004. Estimating equations for association structures. Stat Med 23: 859–880.

Yang, C.C., Wang, L.W., Liang, W. & Moller, A.P. 2017. How cuckoos find and choose host nests for parasitism. Behav Ecol 28: 859–865.

Yom-Tov, Y. 2001. An updated list and some comments on the occurrence of intraspecific nest parasitism in birds. Ibis 143: 133–143

Webster, M. S. 1994. Interspecific brood parasitism of Montezuma oropendolas by giant cowbirds: parasitism or mutualism? Condor: 96, 794–798.

Wood, D.R. & Bollinger, E.K. 1997. Egg removal by brown-headed cowbirds: A field test of the host incubation efficiency hypothesis. Condor 99: 851–857.

Wyllie, I. 1981. The cuckoo. Batsford, London.

